# Thermodynamic constraints on the regulation of metabolic fluxes

**DOI:** 10.1101/341271

**Authors:** Ziwei Dai, Jason W. Locasale

**Author notes:** Correspondence (Z.D.), (J.W.L.).

## Abstract

Nutrition and metabolism are fundamental to cellular function in physiological and pathological contexts. Metabolic activity (i.e. rates, flow, or most commonly referred to as flux) is constrained by thermodynamics and regulated by the activity of enzymes. The general principles that relate biological and physical variables to metabolic control are incompletely understood. Using metabolic control analysis in several representative topological structures of metabolic pathways as models, we derive exact results and conduct computer simulations that define relationships between thermodynamics, enzyme activity, and flux control. We confirm that metabolic pathways that are very far from equilibrium are controlled by the activity of upstream enzymes. However, in general, metabolic pathways have a more adaptable pattern of regulation, controlled minimally by thermodynamics and not necessarily by the specific enzyme that generates the given reaction. These findings show how the control of metabolic pathways, which are rarely very far from equilibrium, is largely set by the overall flux through a pathway rather than by the enzyme which generates the flux or by thermodynamics.

## Introduction

Metabolism enables the utilization of nutritional resources to provide energy, material, and cellular communication for functions of cells(1). Some metabolic pathways such as central carbon metabolism which processes macronutrients or the major caloric sources in diet (proteins, fats, carbohydrates) have been largely defined for over 50 years(2), and even genome-scale reconstructions of metabolic networks which include thousands of metabolites and reactions in different intracellular compartments are available in many unicellular organisms and metazoans(3-5). Nevertheless, the principles that underlie the control of metabolic pathway activity, that is the flow of materials through the network, (i.e. rates, or most commonly referred to as fluxes) are relatively less understood.

Analytical frameworks have been developed to understand steady state and dynamic behaviors of metabolic networks. The most widely applied method in computational modeling of metabolism is flux balance analysis (FBA), which assumes that the network is in steady state and that configurations of metabolic fluxes are determined by optimizing an objective function such as growth rate(6). This approach does not require that enzyme properties such as expression levels are known, but the utility of the method is dependent on the objective function and other assumptions such as nutrient uptake rates. Moreover, FBA does not provide information about pathway regulation and the control of metabolic flux which requires additional knowledge such as enzyme activities or metabolite abundances which are routinely measurable(7,8).

Another framework for understanding metabolism is metabolic control analysis (MCA) originally developed in the 1970s once the biochemistry for many of the key pathways in metabolism such as glycolysis and the tricarboxylic acid (TCA) cycle was established(9-15). MCA quantitatively measures how the flow through a metabolic pathway responds to changes in environmental variables such as the abundance of an enzyme or availability of a nutrient. MCA defines the sensitivity of a metabolic flux to a perturbation in a given metabolic reaction, i.e. the flux control coefficients (FCCs), and also provides a series of rigorously derived relationships between metabolic fluxes, metabolite concentrations, and enzyme activities. This framework can also be applied in computer simulations(16-19) or in experimentation (20,21) when some but not all of the relevant variables are measured(22-25).

All of biology and particularly metabolism are subject to the laws of thermodynamics. These laws place constraints on the dynamics of metabolic reactions(26-32) and are known to affect the control of fluxes in linear pathways(9,31,33,34). Moreover, the deviation from equilibrium at an individual reaction step has been applied as the criterion to identify rate-limiting steps in metabolic pathways(2,29). However, this rule of thumb has been challenged by MCA at least for linear pathways(9), but a quantitative evaluation of the relationship between thermodynamics and flux control in metabolic pathways with different topologies such as the control at branching points is still lacking.

In this study, we use a set of models with representative topological structures observed in metabolism to investigate the quantitative relationships between thermodynamics and regulation of metabolic fluxes. To our surprise, we find that, in both linear and branched pathways, the regulation of pathway fluxes by individual enzymes is in general loosely constrained by the deviation from thermodynamic equilibrium, or the thermodynamic driving force, of the whole pathway. Only pathways very far from equilibrium have their flux regulation strictly constrained by the thermodynamic driving force, in which all fluxes are almost fully controlled by the upstream enzymes. These results unravel simple principles of how metabolic pathways are regulated by the interaction between thermodynamics and enzyme activity.

## Results

### Metabolic flux and thermodynamics

We first consider unimolecular, first-order kinetics of reversible metabolic reactions for simplicity. In this framework, each reaction has one substrate (S) and one product (P) (Fig 1). It is noteworthy that this simplified case approximates the more complicated Michaelis-Menten mechanism when the abundance of substrate is far below the Michaelis-Menten constant, *K_m_*. The forward reaction rate *v*_+_ = *k*_+_*S* and backward reaction rate *v*_−_ = *k*_−_*P* are linear in substrate and product concentrations. Since the rate constants of the forward and backward reactions are coupled by the equilibrium constant 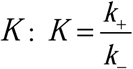, the net flux carried by this reaction is: 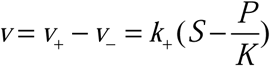. Let *k* = *k*_+_ for simplicity, we have:
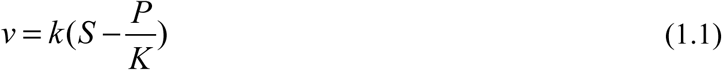

The equilibrium constant *K* can be further connected to the standard Gibbs free energy of this reaction:
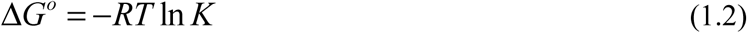

**Figure 1.**
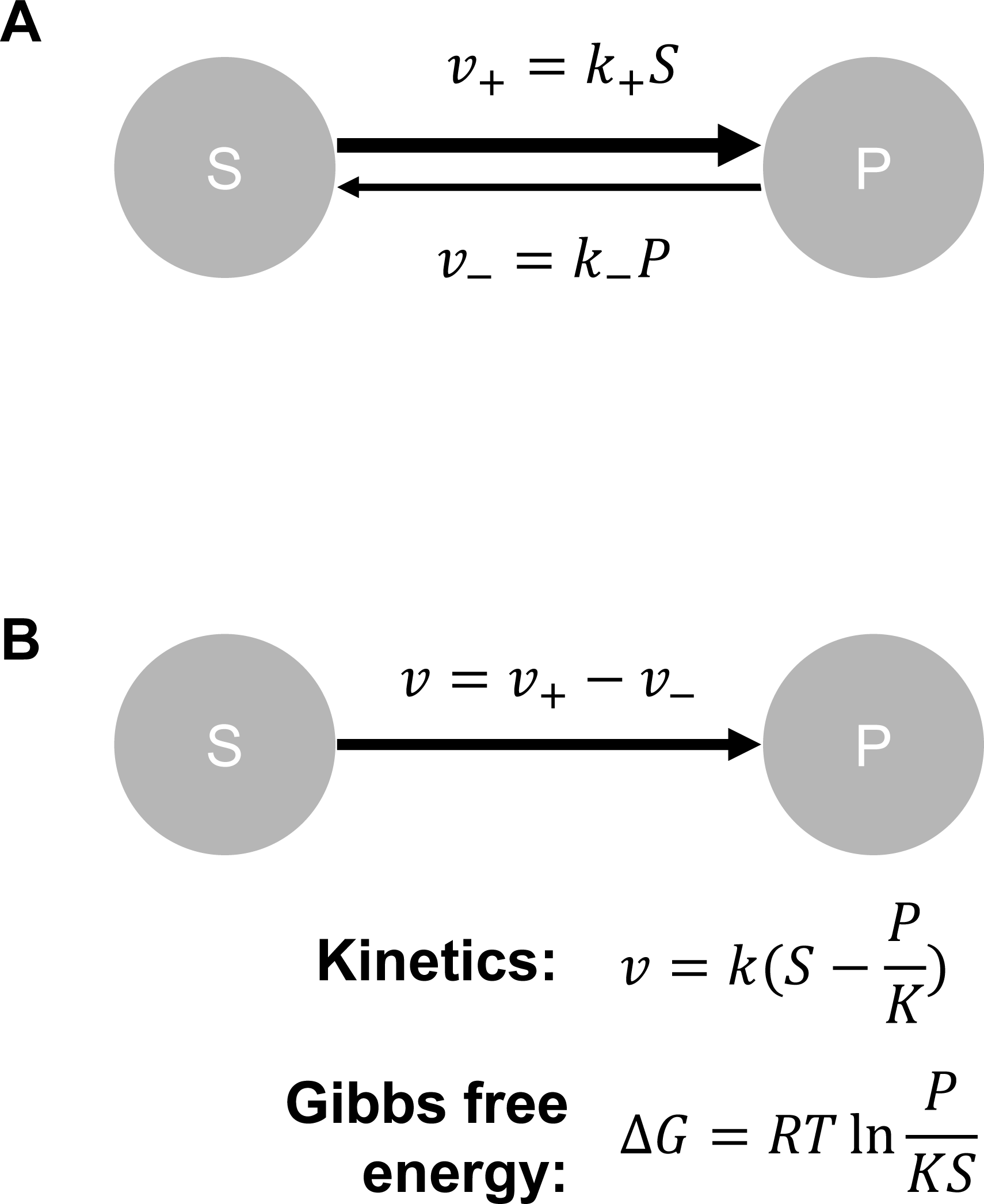
First-order kinetics of enzyme-catalyzed reversible reactions. A. Forward and backward fluxes of a reversible reaction. S is the concentration of substrate, P is the concentration of product, *v*_+_ is the rate of the forward reaction, *v*_−_ is the rate of the backward reaction, *k*_+_ is the rate constant of the forward reaction, *k*_−_ is the rate constant of the backward reaction. B. Net flux and Gibbs free energy change of a reversible reaction. *v* is the net reaction rate, *k* is the rate constant, *K* is the equilibrium constant, Δ*G* is the reaction Gibbs free energy change, *R* is the universal gas constant, *T* is the temperature. Other variables are the same as in (A).

Finally, the Gibbs free energy is determined by the standard Gibbs free energy in combination with concentrations of the substrate and product:
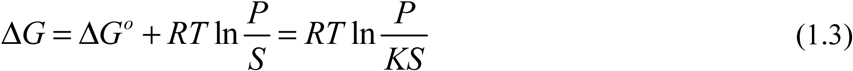

We note that (1.1) can also be rewritten to explicitly incorporate the reaction free energy change. Here we let 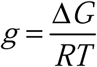 for simplicity. Thus, we have:
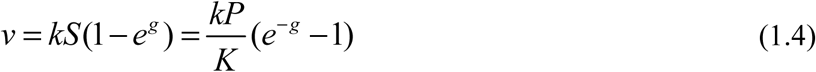

According to the connectivity theorem in MCA(13), flux control coefficients of reactions directly associated with a metabolite are coupled through local elasticity coefficients of these reactions with respect to the metabolite. For a reaction and a metabolite directly associated with this reaction (either as substrate or as product), the elasticity coefficient is defined as the partial derivative of the reaction rate with respect to the concentration of the metabolite on the logarithmic scale. The elasticity coefficients can be computed from (1.1) and (1.4):
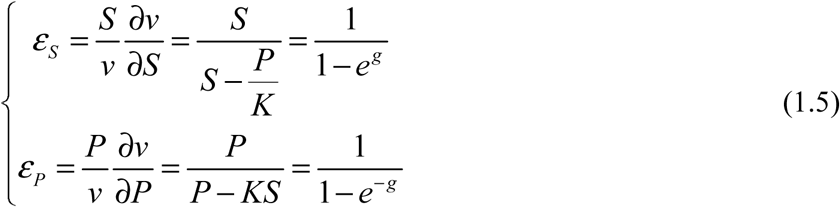

## Linear pathways

Based on the definitions in the previous section, we now consider a linear pathway consisting of *n* reactions (Fig 2A). Each reaction has unimolecular, linear kinetics as described previously. The first substrate *S_in_* is converted to the end product *S_out_* by this reaction chain in *n* steps with *n* − 1 intermediary metabolites *S*_1_,…, *S_n_*_−1_. The *i*-th reaction in this chain has rate constant *k_i_* and equilibrium constant *K_i_*. From MCA(13), we derive analytical relationships between thermodynamic properties and the flux control coefficients, which quantify the relative importance of each enzyme in regulating the flux through the pathway.

**Figure 2.**
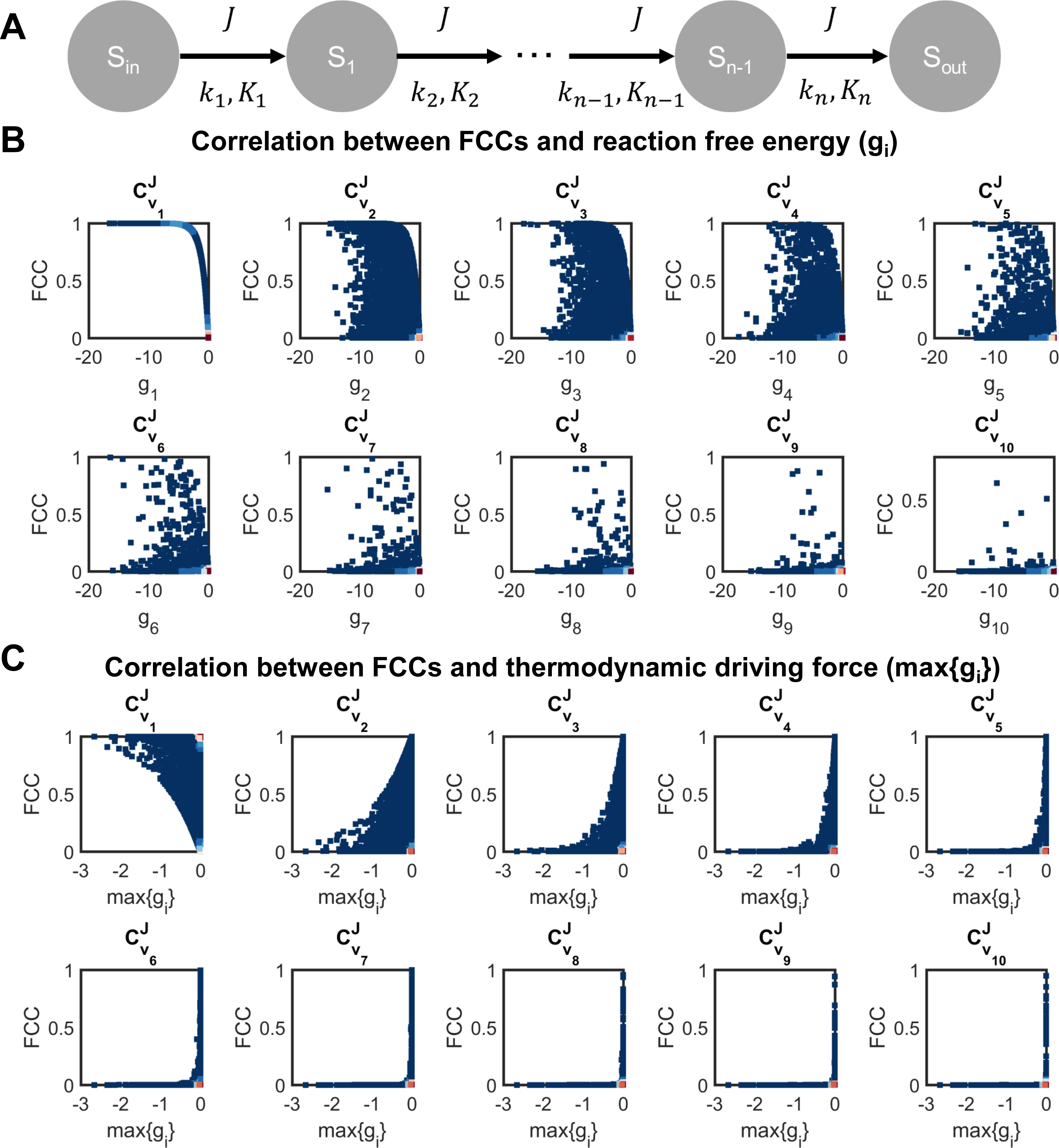
Thermodynamics and flux regulation in a linear pathway. A. Diagram of a linear pathway and related parameters. *S_in_* is the input substrate, *S_out_* is the final product, *S_i_* is the *i* -th intermediary metabolite, *k_i_* is the rate constant of the *i* -th reaction, *K_i_* is the equilibrium constant of the *i* -th reaction, *J* is the pathway flux. B. Scatter plots comparing flux control coefficients and reaction free energy changes in a linear pathway with randomly sampled parameters. The pathway includes 10 reaction steps. 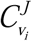 is the *i* -th flux control coefficient and 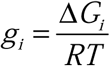 quantifies the *i* -th free energy change. FCC is the abbreviation of flux control coefficient. C. Scatter plots comparing flux control coefficients and the thermodynamic driving force (max{*g_i_*}) in a linear pathway with randomly sampled parameters. Variables are the same as in (B).

To apply the summation theorem and connectivity theorems in the theory of metabolic control analysis, we assume that the pathway is in steady state, in which all reactions have identical net flux. In other words, all intermediary metabolites have balanced fluxes feeding and consuming them. Thus, we can write *n* − 1 equations for the steady state constraints:
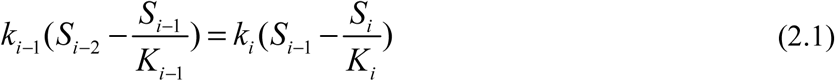

Let *J* denote the net flux through this pathway. The flux control coefficient, 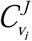, quantifies the sensitivity of the pathway flux *J* to perturbation in activity (i.e. rate constant) of the *i* -th reaction. It is defined as the ratio of relative change in the flux *J* to relative change in the rate of the *i* -th reaction when an arbitrary parameter, *P*, has a small change:
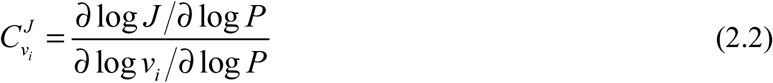

If *P* = *k_i_*, we have:
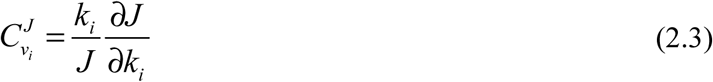

The flux control coefficients can be uniquely determined by solving equations derived from the summation and connectivity theorems. For linear pathways, the summation and connectivity theorems don’t explicitly include the pathway flux *J*:

<u>Summation theorem</u>
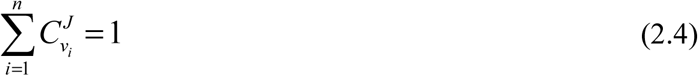

<u>Connectivity theorem</u>
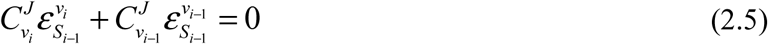

According to (1.5), we have:
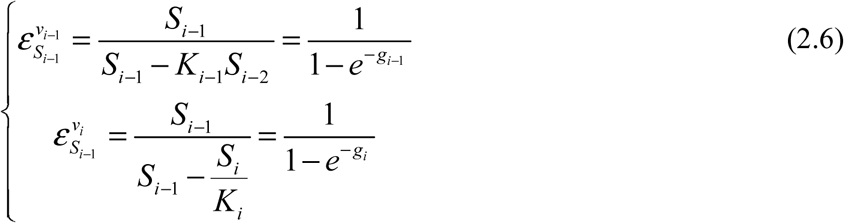

Flux control coefficients can be solved by combining (2.1), (2.4), (2.5) and (2.6). They can be written as either explicitly including only the reaction free energy terms or explicitly including only the rate constants and equilibrium constants:
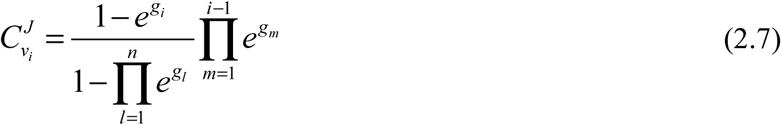

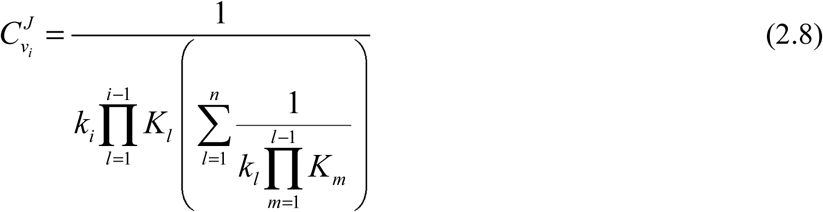

According to (2.7) and (2.8), the flux control coefficients are completely determined by kinetic and thermodynamic parameters of reactions in the pathway. Therefore, alterations in concentrations of the input substrate and the final product don’t influence the flux control coefficients.

To illustrate the quantitative relationships defined in (2.7) and (2.8), we consider a linear pathway consisting of 10 reactions, randomly sample 20,000 combinations of the parameters *S_in_*, *S_out_*, {*k_i_*} and {*K_i_*}, and compute the corresponding flux control coefficients. We select the size 10 because it approximates the typical length of linear metabolic pathways. All parameters and substrate concentrations are sampled from a log-normal distribution:
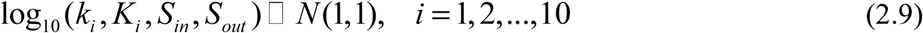

We first correlate the flux control coefficient with reaction free energy change for each individual reaction (Fig 2B). We find that the relationship between reaction free energy changes and flux control coefficients questions the long-standing hypothesis that reactions with most negative free energy changes, such as the reactions catalyzed by hexokinase, phosphofructokinase and pyruvate kinase in glycolysis, serve as rate-limiting steps of a pathway(35,36). For all reactions except the first reaction, their flux control coefficients correlate poorly with the reaction free energy changes, suggesting that regulation of metabolic fluxes by an individual reaction is determined by both thermodynamics property and position along the pathway.

We next investigate how the flux control coefficients are associated with global thermodynamic properties of the entire pathway. We quantify the deviation from the thermodynamic equilibrium by the reaction free energy change that is closest to zero (i.e. the maximal value of free energy change among all reactions). The pathway has zero net flux (i.e. thermodynamic equilibrium) when this quantity equals to zero. Thus, it is termed the thermodynamic driving force (TDF)(34):
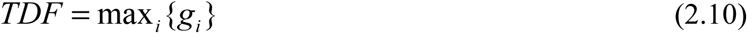

We correlate the thermodynamic driving force with flux control coefficient of each reaction step (Fig 2C). From these simulations, a pattern emerges, in which the thermodynamic driving force determines either upper bound or lower bound of the flux control coefficients: for the flux control coefficient of the first reaction 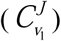, its lower bound is dependent on the thermodynamic driving force; while for the rest reactions, the upper bounds of their flux control coefficients depend on the thermodynamic driving force.

Analytical relationships that constrain flux control coefficients by thermodynamic driving force can also be derived from (2.7) and (2.10) (Supplementary Information):
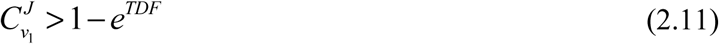

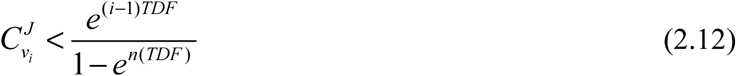

From (2.11) and (2.12), we can derive corollary relationships for the flux control coefficients when the system is very far from equilibrium (i.e. thermodynamic driving force approaches minus infinity):
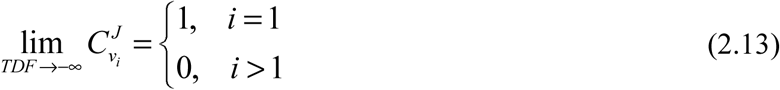

Equation (2.13) suggests that in a linear pathway far from thermodynamic equilibrium, the metabolic flux through the pathway is fully controlled by the enzyme catalyzing the first step.

To summarize, for a linear metabolic pathway, we have derived analytical relationships between the flux control coefficients and thermodynamic properties of both individual reactions and the entire pathway. We have shown that the flux control coefficients correlate poorly with free energy changes of the corresponding reactions but that the thermodynamic driving force of the whole pathway places bound on how much any enzyme can control pathway flux. However, when the pathway is very far away from thermodynamic equilibrium, the metabolic flux through this pathway is completely controlled by the first reaction. We next investigate if these principles are conserved in metabolic networks with more complex topologies.

## Branch point with two upstream fluxes

We next consider a metabolic network with two converging fluxes, *J*_1_ and *J*_2_, at a branch point (Fig 3A). Through the two fluxes, the metabolite at the branch point, *S_BP_*, is produced by two input substrates, *S_in_*_,1_ and *S_in_*_,2_. The final product *S_out_* is produced from *S_BP_* with the flux *J*_1_ + *J*_2_ at the steady state. The three reactions included in this network have rate constants {*k*_1_, *k*_2_, *k*_3_} and equilibrium constants {*K*_1_, *K*_2_, *K*_3_}. At the steady state, the fluxes *J*_1_ and *J*_2_, and the concentration of *S_BP_* can be solved from the rate constants, equilibrium constants, and concentrations of the input substrates and end product based on the kinetic rules:
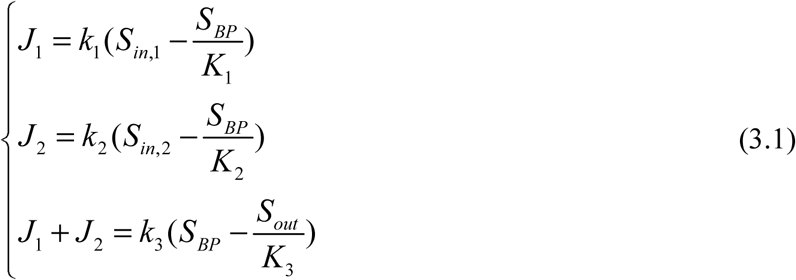

From (3.1) we can solve the steady state concentration of the branch point metabolite, *S_BP_*:
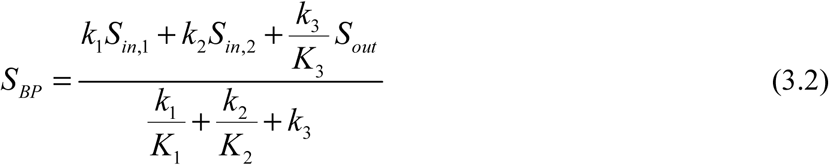

**Figure 3.**
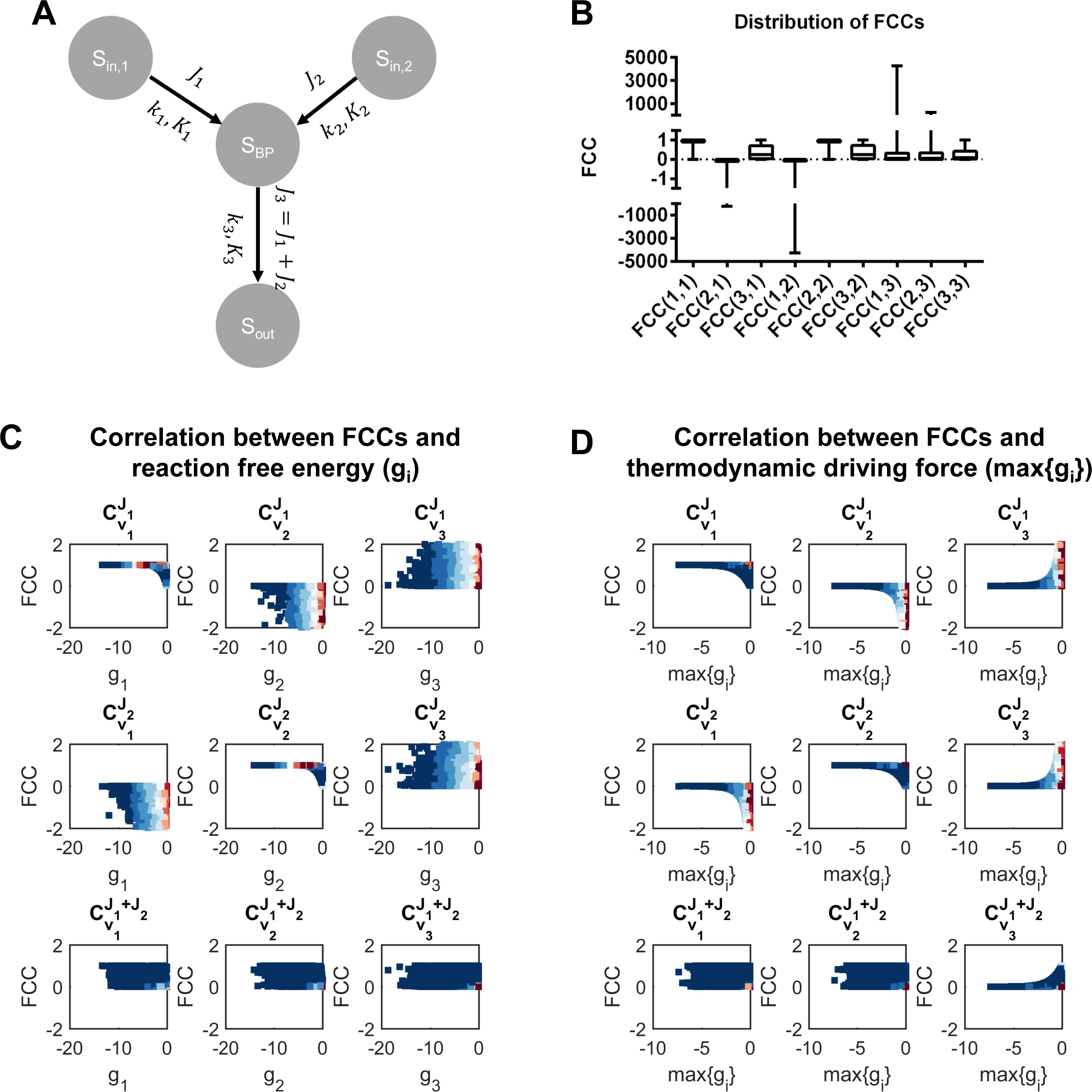
Thermodynamics and flux regulation in a pathway with a branch point and two upstream fluxes. A. Diagram of the pathway and related parameters. *S_in_*_,1_ and *S_in_*_,2_ are the two input substrates, *S_out_* is the final product, *S_BP_* is the intermediary metabolite at branch point, *k_i_* is the rate constant of the *i* -th reaction, *K_i_* is the equilibrium constant of the *i* -th reaction, *J*_1_ and *J*_2_ are two upstream fluxes, *J*_3_ = *J*_1_ + *J*_2_ is the downstream flux. B. Distributions of the flux control coefficients for the network in (A) with randomly sampled parameters. 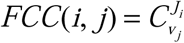 is the flux control coefficient of the *i* -th flux with respect to the *j* -th reaction. Limits of the boxes are the 25^th^ and 75^th^ percentiles, central lines are median values, whiskers indicate minimal and maximal values. C. Scatter plots comparing flux control coefficients and reaction free energy changes in the network in (A) with randomly sampled parameters. 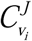 is the *i* -th flux control coefficient and 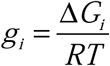 quantifies the *i* -th free energy change. D. Scatter plots comparing flux control coefficients and the thermodynamic driving force (max {*g_i_*}) in the network in (A) with randomly sampled parameters. Variables are the same as in (C).

And the steady state fluxes:
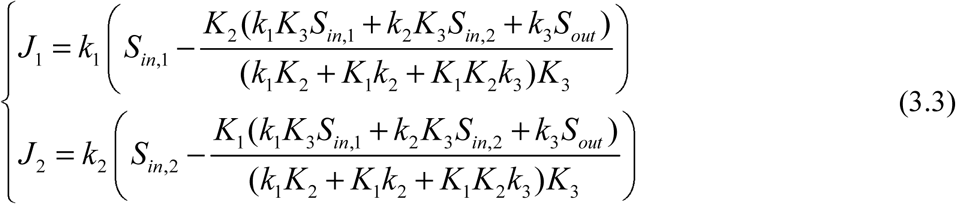

We can then determine the flux control coefficients from the summation and the connectivity theorems. Here we have three fluxes and three reactions in the network, giving 9 flux control coefficients:
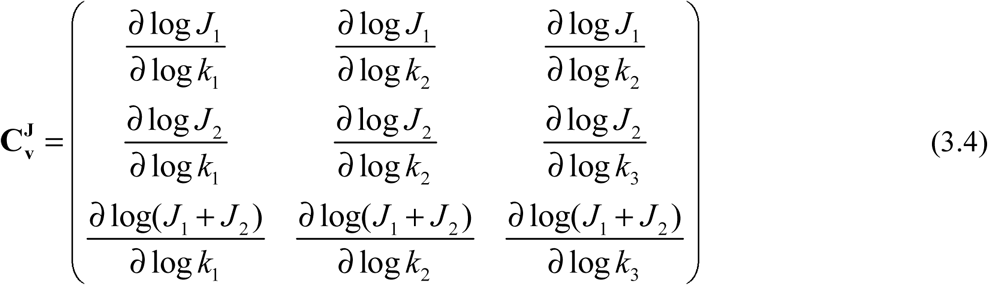

These flux control coefficients can be determined based on the summation theorem and connectivity theorem:

<u>Summation theorem</u>
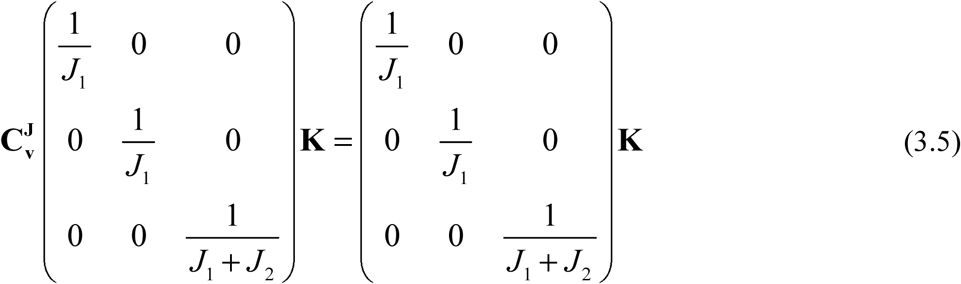

In which columns of **K** are the linear basis of feasible steady state flux configurations in the network (i.e. all feasible steady state flux configurations can be written in the form of linear combination of columns of **K**):
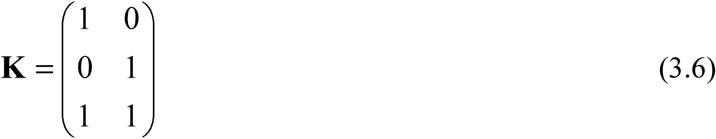

<u>Connectivity theorem</u>
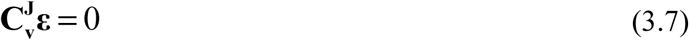

In which **ε** is the matrix consisting of the normalized elasticity coefficients:
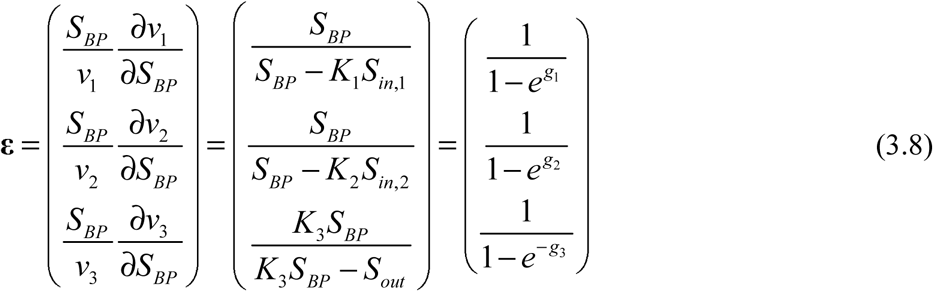

Thus, by combining (3.1)-(3.8), the flux control coefficients can be uniquely solved:
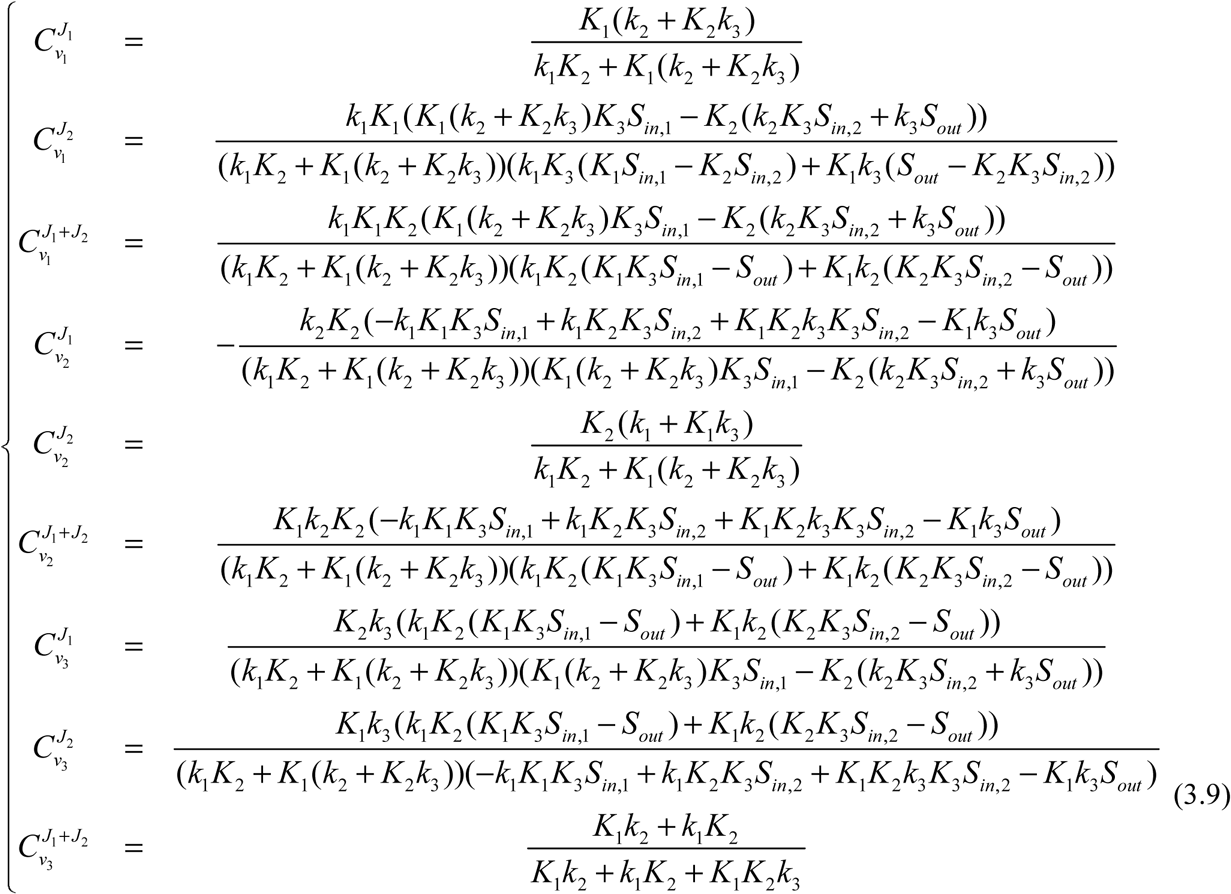

Furthermore, the reaction free energy changes can also be written as functions of the kinetic parameters and substrate concentrations:
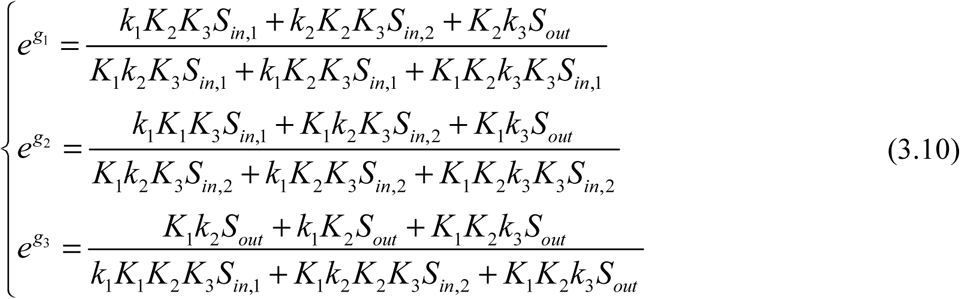

We then combine (3.9) and (3.10) to express the flux control coefficients in terms of the reaction parameters and free energy terms. To simplify the expressions, we let 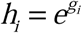. Then we have:
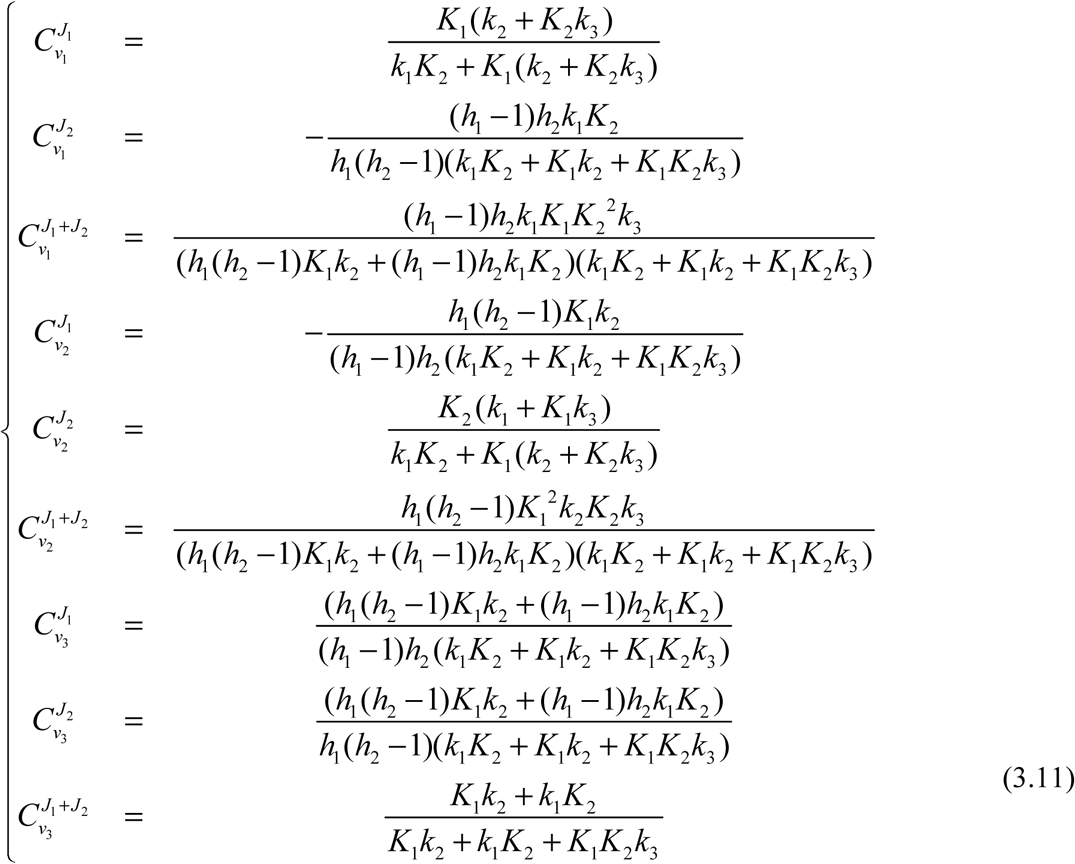

To illustrate the relationships between flux control coefficients and thermodynamic properties of the network, according to (3.9) - (3.11), we simulate the flux control coefficients and reaction free energy changes based on 20,000 combinations of parameters randomly sampled from a log-normal distribution as previously described. Combinations of parameters resulting in negative fluxes are discarded. The distributions of flux control coefficients (Fig 3B) show that 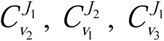 and 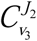 can have absolute values much larger than 1 under certain parameter combinations, indicating that the two upstream fluxes can be dramatically altered in response to perturbation of activities of enzymes in other branches. Based on the simulation, we also compare flux control coefficients with the reaction free energy changes (Fig 3C) and the thermodynamic driving force (Fig 3D). Consistent with the case of the linear pathway, most of the flux control coefficients correlate poorly with free energy changes of the individual reactions catalyzed by the corresponding enzyme (Fig 3C) but exhibit a dependence on the thermodynamic driving force (Fig 3D). Moreover, the quantitative relationships characterizing how flux control coefficients are constrained by the thermodynamic driving force can also be analytically derived (Supplementary Information):
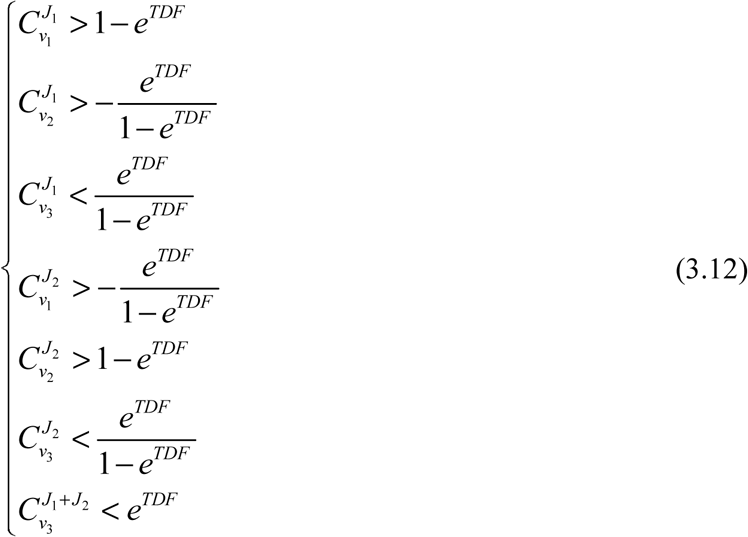

From (3.12), we can derive the limits of all the flux control coefficients except for 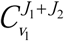 and 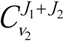 when the thermodynamic driving force goes to minus infinity:
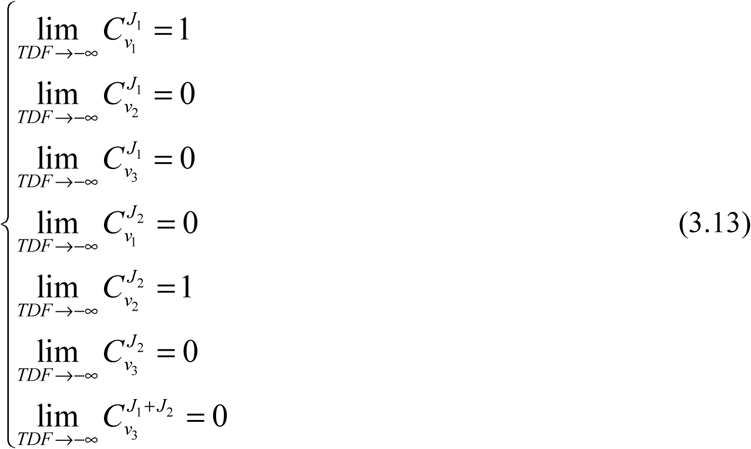

Thus, when the pathway is far from equilibrium, the two upstream fluxes *J*_1_ and *J*_2_ are fully regulated by activities of the enzymes that generate them. Moreover, the downstream flux *J*_1_ + *J*_2_ is also fully controlled by the two upstream enzymes. The inequalities in (3.12) also indicate that the feasible regions of the flux control coefficients 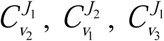 and 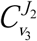 become semi-infinite (i.e. only bounded in one direction) when the thermodynamic driving force approaches zero (i.e. there exists at least one near-equilibrium reaction in this pathway).

To summarize, in a metabolic pathway with a branch point involving two converging fluxes, regulation of the fluxes is largely dependent on how close to thermodynamic equilibrium the pathway in entirety is. When the pathway is far away from equilibrium, all three fluxes are completely controlled by the two upstream enzymes. On the other hand, when there exists at least one near-equilibrium reaction in the pathway, there is more flexibility in the regulation of fluxes, and the two upstream fluxes are able to be dramatically altered even by enzymes that aren’t directly involved in the reaction.

## Branch point with two downstream fluxes

We next consider a pathway with one branch point in which one upstream flux, *J*_1_, diverges to two downstream fluxes *J*_2_ and *J*_3_, and show that the principles of metabolic flux regulation and thermodynamics also hold in this case. This network includes one input substrate, *S_in_*, and two final products, *S_out_*_,1_ and *S_out_*_,2_, which are all connected to the branch point metabolite *S_BP_*. All kinetic rules and assumptions are as in the previous sections. Thus, at the steady state, concentration of *S_BP_* can be solved from the reaction parameters and concentrations of other metabolites in the network:
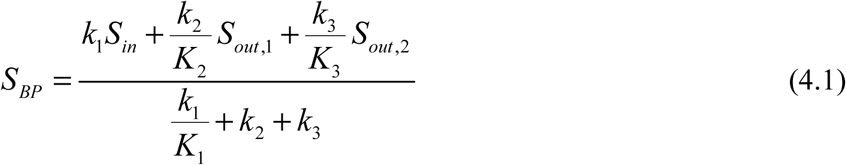

Fluxes and reaction free energy changes at the steady state can also be calculated from (4.1):
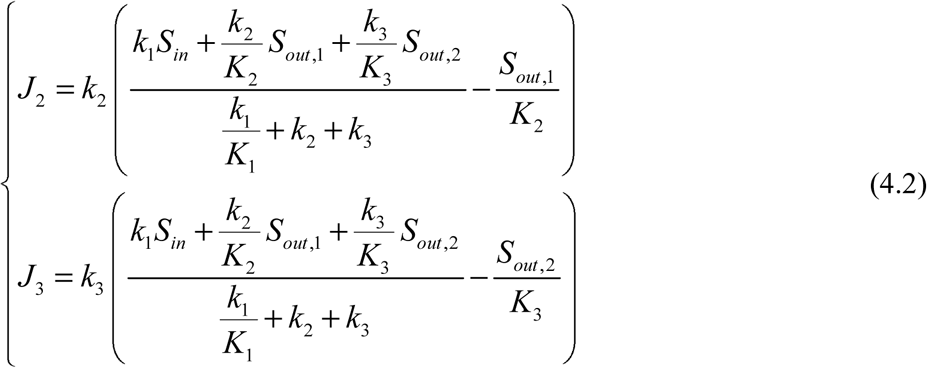

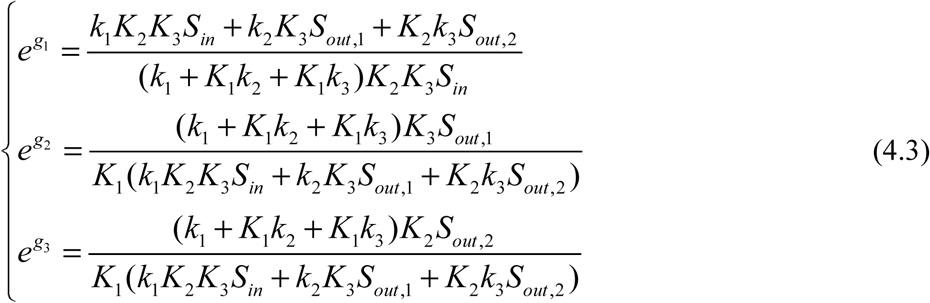

Here we also have in total nine flux control coefficients which can be solved based on the summation theorem and connectivity theorem:
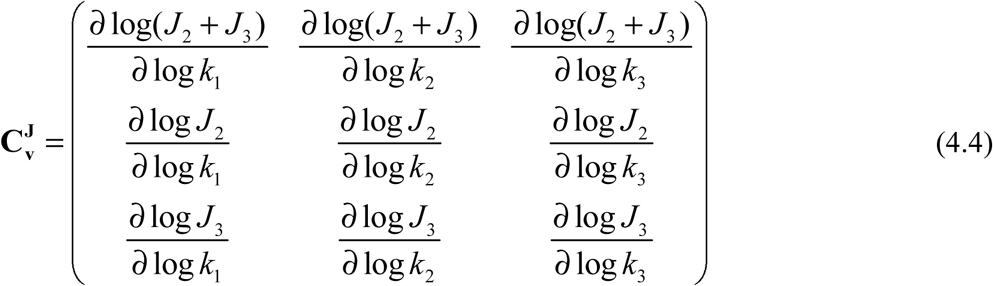

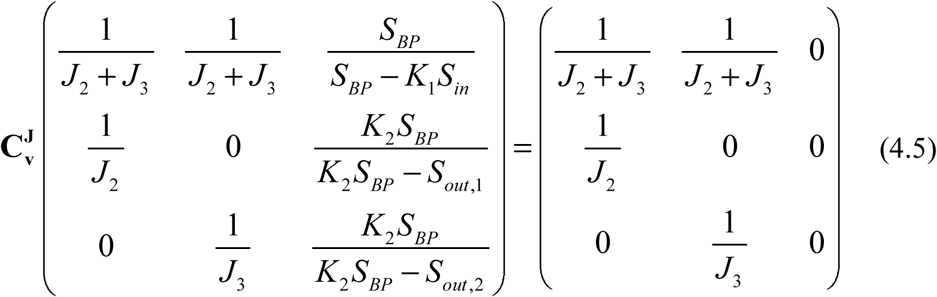

By combining (4.2)-(4.5), we can also determine the flux control coefficients as functions of reaction parameters and free energy changes:
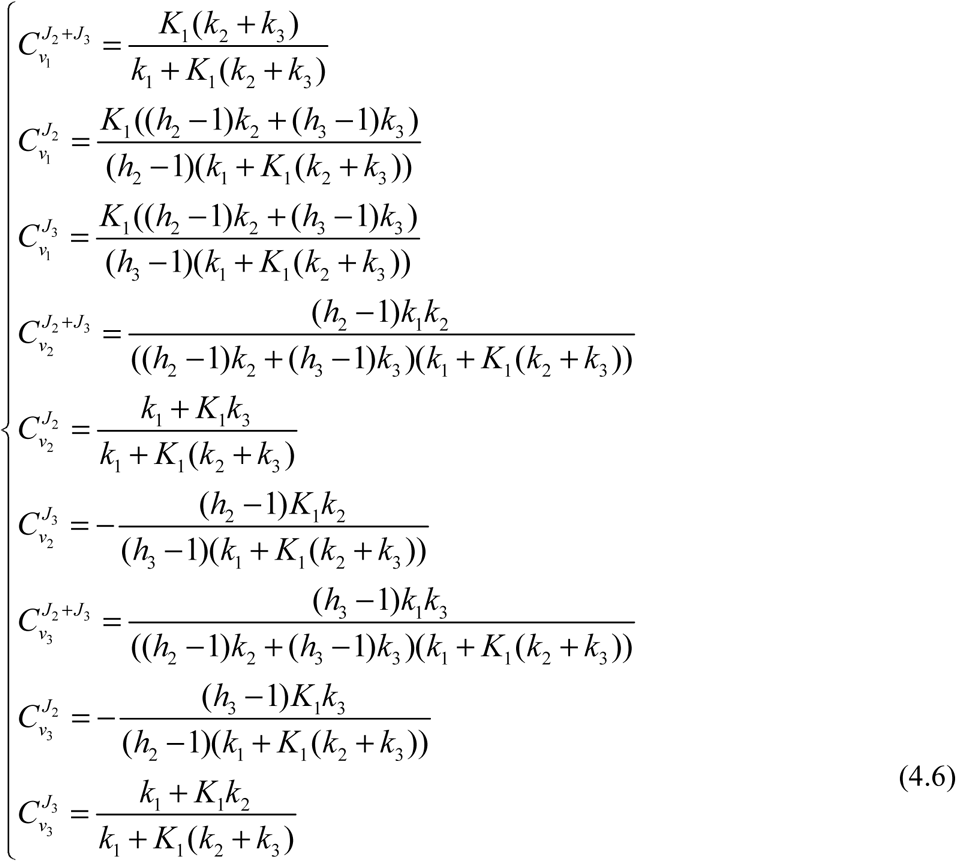

To investigate the relationships between flux control coefficients and thermodynamic variables, we randomly sample 20,000 combinations of reaction parameters and substrate concentrations and calculate flux control coefficients and free energy changes corresponding to the randomly sampled parameters from (4.3) and (4.6). Distributions of the sampled flux control coefficients suggest that the two downstream fluxes are able to be regulated in response to variations in the activity of enzymes not directly carrying them (i.e. 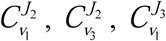 and 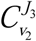 can have absolute values larger than 1, Fig 4B). Again, we observe a poor correlation between flux control coefficients and free energy change of the corresponding reaction steps (Fig 4C). Among the 9 flux control coefficients, only 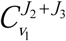 exhibits a clear dependence on the free energy change *g*_1_.

**Figure 4.**
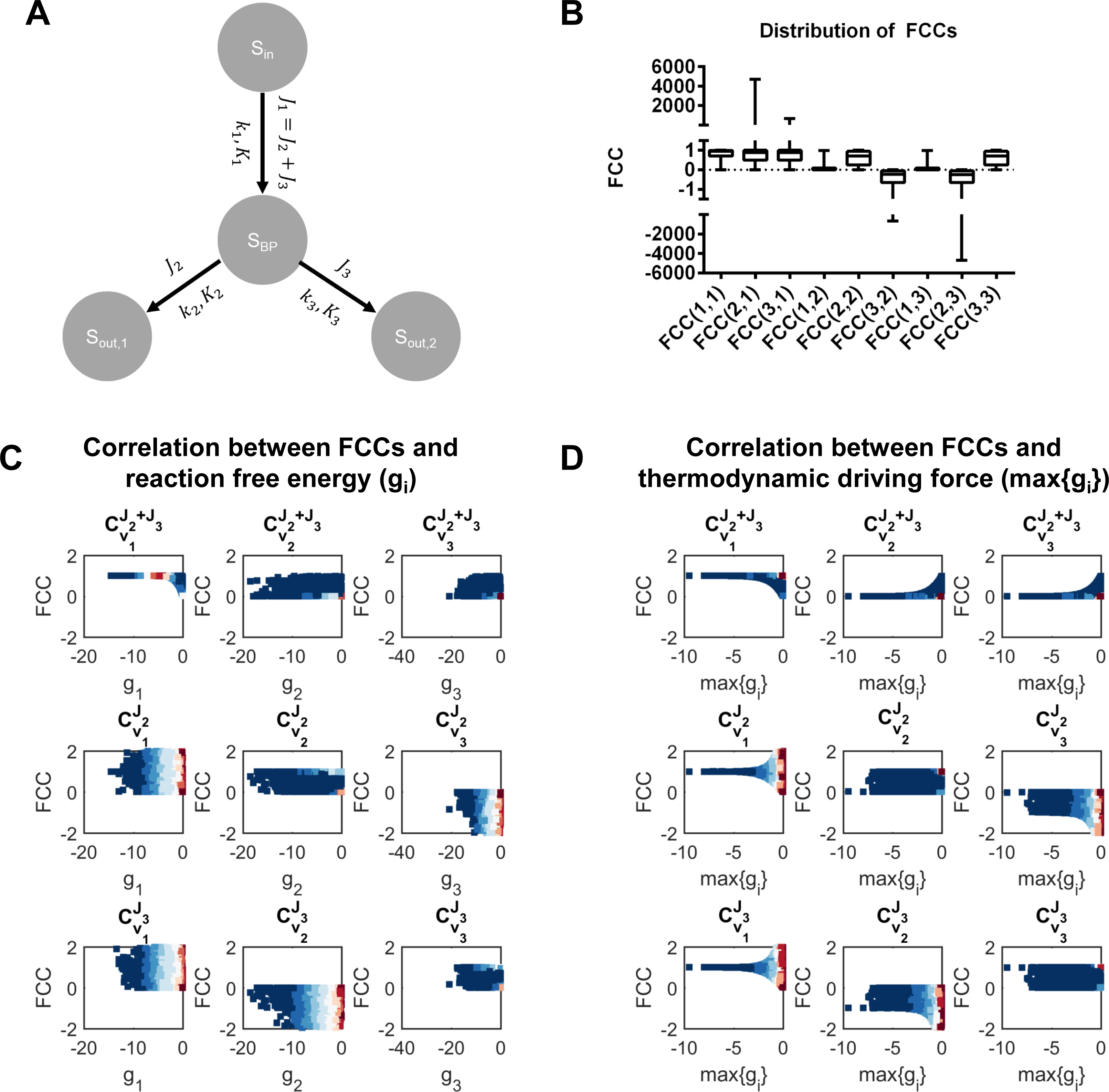
Thermodynamics and flux regulation in a pathway with a branch point and two downstream fluxes. A. Diagram of the pathway and related parameters. *S_in_* is the input substrate, *S_out_*_,1_ and *S_out_*_,2_ are the two final products, *S_BP_* is the intermediary metabolite at branch point, *k_i_* is the rate constant of the *i* -th reaction, *K_i_* is the equilibrium constant of the *i* -th reaction, *J*_2_ and *J*_3_ are two downstream fluxes, *J*_1_ = *J*_2_ +*J*_3_ is the upstream flux. B. Distribution of the flux control coefficients for the network in (A) with randomly sampled parameters. 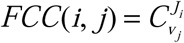 is the flux control coefficient of the *i* -th flux with respect to the *j-* th reaction. Limits of the boxes are the 25^th^ and 75^th^ percentiles, central lines are median values, whiskers indicate minimal and maximal values. C. Scatter plots comparing flux control coefficients and reaction free energy changes in the network in (A) with randomly sampled parameters. 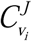 is the *i* -th flux control coefficient and 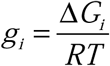 quantifies the *i* -th free energy change. D. Scatter plots comparing flux control coefficients and the thermodynamic driving force (max {*g_i_*}) in the network in (A) with randomly sampled parameters. Variables are the same as in (C).

Despite the poor correlation between flux control coefficients and free energy changes of individual reaction steps, 7 of the 9 flux control coefficients are strictly constrained by the thermodynamic driving force (Fig 4D). This corroborates the findings in other types of network topologies that the regulation of metabolic fluxes is constrained by the deviation of the whole pathway from thermodynamic equilibrium. We also analytically derive the quantitative relationships between the upper and lower bounds of the flux control coefficients and the thermodynamic driving force of the pathway (Supplementary Information):
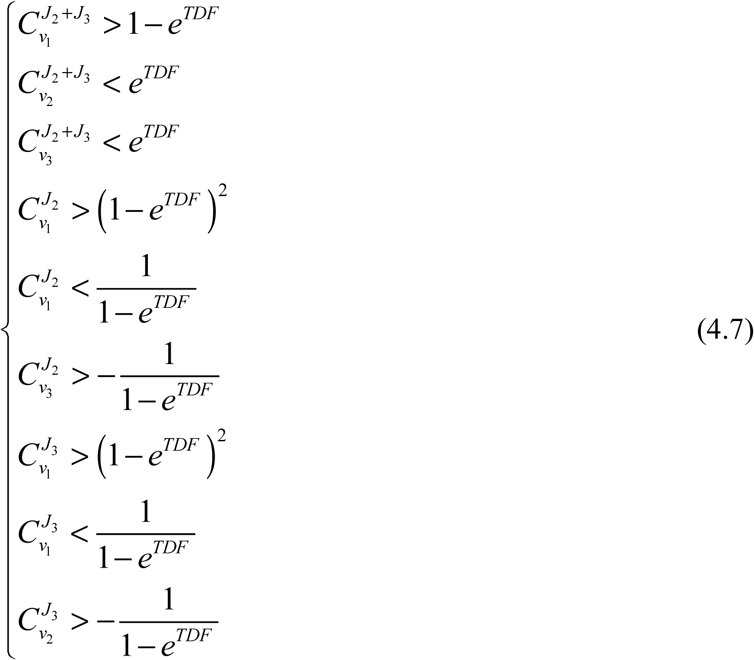

According to (4.7), when the network moves away from the thermodynamic equilibrium, many of the flux control coefficients converge to fixed values:
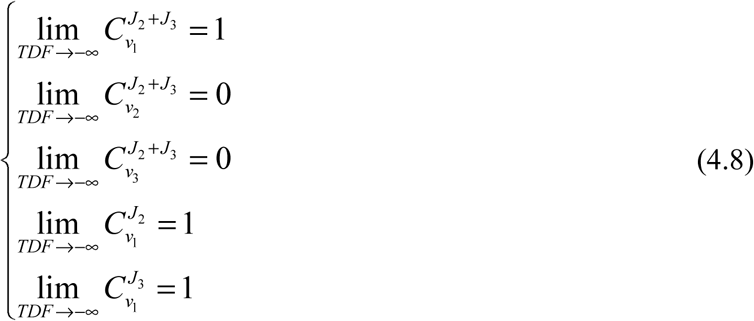

Thus, when the pathway is far away from thermodynamic equilibrium, the enzyme catalyzing the upstream reaction exerts complete control of all three fluxes. Larger variability in all flux control coefficients that result in semi-infinite feasible regions of the flux control coefficients 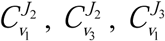 and 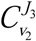 are allowed only when there exists near-equilibrium reactions in this network.

Thus, based on a combination of analysis and simulation, we have demonstrated that, at branch points, the regulation of fluxes by enzyme activities is constrained by the thermodynamic driving force but has little correlation with free energy changes of the individual enzymes. These findings, together with similar results in other network structures, suggest that the influences of thermodynamics on regulation of metabolic fluxes by enzyme activities primarily occur at the pathway level rather than at the individual reaction level.

## Discussion

In this study, we derive quantitative relationships between thermodynamics, enzyme activity and regulation of metabolic fluxes. For a set of example pathways, we calculate the flux control coefficients as functions of the enzyme rate constants, reaction equilibrium constants and Gibbs free energy. Based on numerical simulation and exact analytical results, we find that in all network topologies considered, the flux control coefficients are bounded by the thermodynamic driving force, as defined by the deviation of the entire pathway from thermodynamic equilibrium. Moreover, distinct patterns of flux regulation emerge when the thermodynamic driving force approaches two extreme values: if the thermodynamic driving force is very negative (i.e. all reactions are far away from equilibrium), enzymes catalyzing the upstream reactions (i.e. reactions directly consume the input substrates) exert full control of all fluxes; on the other hand, if the thermodynamic driving force is close to zero (i.e. near-equilibrium reactions exist in the pathway), there is more flexibility in the pattern of flux control, and regulation of fluxes in network topologies with branch points (i.e. switching from one branch point to another) is allowed. These findings suggest that the coupling between thermodynamics and regulation of metabolic flux occurs at the systemic level and challenges the rule of thumb that the reaction with the most negative free energy change serves as the rate-limiting step.

It is also worth noting that all analysis that we have done here relies on the assumption of first-order kinetics. This approximates the more generalized Michaelis-Menten mechanism when the abundance of substrate is far below the K_m_ of the enzyme. Accordingly a study comparing substrate concentrations and K_m_ values in different cell types, among all substrate-enzyme interactions investigated, around 70% exhibited higher substrate concentration than the K_m_ value, thus requiring a Michaelis-Menten or substrate-saturated, zero-order kinetics(30). We also note that the tendency of an enzyme to be operating in the linear region or to be saturated by its substrate is highly dependent on the substrate used by this enzyme. For instance, kinases use ATP as the substrate whose physiological concentration is much higher than the typical K_m_ value of enzymes, thus are more likely to be substrate-saturated. On the other hand, other enzymes such as chromatin-modifying enzymes use metabolites much less abundant as substrates(37). Nevertheless, our models allowed for exact results and likely will have some generalizations to realizable pathways.

To characterize the relationships between thermodynamics and flux control by substrate-saturated enzymes, we repeated the theoretical analysis and simulation in the same metabolic networks with zero-order kinetics (Supplementary Information). Briefly, although the expressions of steady state concentrations of the intermediary metabolites and steady state fluxes are much more complicated with such assumption, the flux control coefficients can still be written in the form of simple functions of the reaction parameters and free energy changes. However, the thermodynamic driving force no longer constrains the flux control coefficients in this case (Supplementary Fig 1, 2). Most of the flux control coefficients in all three network structures have the form of a zero-order homogenous function of the thermodynamic terms *h_i_*, thus being invariant while simultaneously increasing or decreasing the values of all *h_i_* by multiplying the same factor. Consequently, moving all reactions away from or towards the thermodynamic equilibrium without influencing the ratios between the *h_i_* s doesn’t change the resulting flux control coefficients. Nevertheless, we expect that the thermodynamic driving force still constrains the flux control coefficients in real biochemical reactions since Michaelis-Menten mechanisms lie in between the two limits of first-order and substrate-saturated kinetics.

Finally, although the network structures we study here are much simpler than real metabolic networks, the conclusions we drive here can still be extended to more complicated network topologies. A metabolic network without cycles can always be simplified to a set of branch points connected by linear reaction chains with varying lengths, and the enzymes in the same linear reaction chain can be treated as an entirety to calculate the flux control coefficients with respect to simultaneous change in these enzymes(38). In this case, each linear reaction chain is simplified to a single reaction step, which enables us to apply the principles we found in the simple network structures with a branch point.

To summarize, we characterize the quantitative relationship between thermodynamics and regulation of metabolic fluxes by enzymes in a metabolic network in this study. We find that the global thermodynamic property of the network, i.e. thermodynamic driving force, constrains almost all flux control coefficients in both linear and branched pathways. Specifically, fluxes in metabolic networks far away from thermodynamic equilibrium are almost fully controlled by enzymes catalyzing the upstream reactions (i.e. reactions directly consuming the input substrates). On the other hand, near-equilibrium metabolic networks allow more flexibility in the pattern of regulation. Semi-infinite feasible regions of flux control coefficients are only allowed in branched pathways with near-equilibrium reactions. These findings highlight the importance of global thermodynamic feature in constraining the pattern of regulation of metabolism.

## Methods

Analytical expressions of flux control coefficients were calculated using the function LinearSolve[] in Wolfram Mathematica 11. For the computer simulation, 20,000 combinations of random parameters were generated using the function normrnd() in MATLAB R2016b for each model. Flux control coefficients and reaction free energies were then computed using the parameter sets that generate positive flux configurations. Source codes are available at GitHub page of the Locasale Lab <u>(https://github.com/LocasaleLab/MCA_thermodynamics).</u>

## Acknowledgements

Support from the National Institutes of Health (R01CA193256, R00CA168997), and the American Cancer Society (TBE-TBE434120) to JWL are gratefully acknowledged. We thank members of the Locasale Lab for helpful discussions.

## Conflict of interest

The authors declare that they have no conflicts of interest with the contents of this article.

